# TubercuProbe: A Cross-Attention Graph-Sequence Model for Cross-Species Chemoproteomic Discovery in *Mycobacterium tuberculosis*

**DOI:** 10.64898/2026.01.04.697588

**Authors:** Abhiram Chalamalasetty, Adesh Rohan Mishra, Fleur M. Ferguson, Benjamin Sanchez-Lengeling, José Manuel Barraza-Chavez, Adrian Jinich

## Abstract

Activity-based protein profiling (ABPP) and residue-specific chemoproteomics have transformed human chemical biology, yet applying these approaches to pathogens remains limited by biosafety constraints, low throughput, and the absence of reusable atlases. We present TubercuProbe, a cross-species machine learning framework that leverages large-scale human chemoproteomic knowledge to prioritize compound-protein interactions in *Mycobacterium tuberculosis* (Mtb) and other pathogens. Our model integrates a graph isomorphism network (GINE) for ligand encoding with frozen ESM-C (600M) protein embeddings via bidirectional cross-attention. Trained on *>*2M ChEMBL compound-protein pairs (predominantly human targets), TubercuProbe achieves *R*^2^=0.77 (MSE=0.45) for continuous affinity prediction and transfers effectively to binary cysteine reactivity prediction (CysDB AUPRC=0.63). Ablation studies reveal that pretrained features are highly transferable across freezing strategies (ΔAUPRC*<*0.03), suggesting the model captures fundamental protein-molecule interaction patterns. As a case study, we prioritize cysteine-reactive electrophiles and molecular glues for three Mtb virulence proteins (PtpB, SapM, Rv3671c), providing candidate probes for prospective ABPP validation. Orthogonal comparison with Boltz-2 structure predictions shows moderate correlation (Pearson *r*≈0.69). TubercuProbe provides a lightweight, sequence-driven first-pass ranker that enables pre-experimental prioritization—reducing the time, cost, and experimental burden of chemoproteomic discovery in biosafety-restricted systems. We discuss extensions toward multitask learning that jointly predicts non-covalent binding and covalent reactivity, recognizing that effective covalent probes must both *reach* their target and *react* once there.

## 1 Introduction

### 1.1 The Translational Gap in Pathogen Chemoproteomics

Activity-based protein profiling (ABPP) and residue-specific chemoproteomics have reshaped drug discovery by enabling systematic mapping of proteome-wide ligandability. In human systems, large-scale efforts such as CysDB and DrugMap have quantified tens of thousands of reactive sites, creating reusable atlases that support translational chemical biology[1]. These advances have fundamentally altered how ligandability is defined and exploited in therapeutic development.

In pathogenic organisms, however, chemoproteomic discovery has not followed the same trajectory. Although ABPP is technically feasible in bacteria, published studies are typically low-throughput, probe-limited, and fragmented—rarely exceeding a few thousand quantified sites per organism. This disparity reflects a convergence of constraints: permeability barriers, efflux mechanisms, slow growth and biomass requirements, biosafety-driven sample inactivation, and the absence of standardized pathogen chemoproteomic resources. As a result, pathogen chemoproteomics remains fragmented across individual studies, preventing cumulative discovery.

### 1.2 Cross-Species Transfer Learning

We hypothesize that machine learning models trained on large-scale human chemoproteomic data can predict ligandable sites across non-human proteomes, including pathogenic bacteria. This cross-species transfer learning strategy leverages conserved biochemical reactivity signatures to translate insights from data-rich human proteomes into biosafety-restricted or data-scarce microbial systems.

Here, we present TubercuProbe, a cross-attention graph-sequence model that addresses this translational bottleneck. Our contributions are:

1. **Cross-attention architecture**: We integrate a graph isomorphism network (GINE) for molecular graphs with frozen ESM-C protein embeddings through bidirectional cross-attention, enabling efficient fusion of molecular and protein representations.
2. **Large-scale pretraining**: Training on *>*2M ChEMBL pairs (predominantly human targets) yields *R*^2^=0.77, capturing generalizable binding patterns that transfer across species.
3. **Transfer to covalent reactivity**: Fine-tuning on CysDB achieves AUPRC=0.63 for cysteine reactivity prediction, with ablation studies revealing high transferability of pretrained features.
4. **Case study in tuberculosis**: We prioritize cysteine-reactive electrophiles and molecular glues for three Mtb virulence proteins (PtpB, Rv3671c, SapM), providing candidates for prospective ABPP validation[2].

Together, these results demonstrate that a scalable, sequence-based model can enable pre-experimental prioritization for pathogen chemoproteomic campaigns, complementing more specialized structural pipelines.

## 2 Related Works

### Sequence/SMILES-only DTA

Early deep models such as DeepDTA learned representations directly from 1D protein sequences and SMILES via CNNs, demonstrating that sequence-only features can predict continuous binding affinities on DAVIS and KIBA benchmarks[3].

### Graph-based ligand + sequence

GraphDTA replaced 1D drug strings with molecular graphs and GNN encoders (GCN, GAT, GIN), paired with 1D protein sequences fed to CNNs, reporting consistent gains over sequence-only baselines[4]. Our ligand encoder follows this lineage but uses GINE (edge-aware GIN) with residual connections, utilizes bidirectional cross-attention, and fuses with frozen ESM-C protein embeddings instead of training protein CNNs, yielding simpler training, higher accuracy, and better generalization[5, 6].

### Structure-aware co-folding

Boltz-1/2 jointly model protein-ligand complexes; Boltz-2 adds an affinity head and reports correlations approaching FEP while remaining far cheaper than physics-based methods[7, 8]. These models deliver stronger absolute accuracy and structures, but at higher per-sample cost.

### Parallels to SimpleFold

SimpleFold replaces domain-specific geometric blocks with general-purpose Transformers trained with flow matching, achieving competitive folding accuracy with reduced complexity[9]. Our philosophy is analogous: trade co-folding for frozen PLM embeddings + GNNs to gain throughput for proteome-scale ranking while displaying significant improvement on intrinsically disordered proteins.

## 3 Methods

### 3.1 Data

We trained the model as a supervised regression task using compound-protein affinity pairs from ChEMBL[10] (labels: pChEMBL = − log^10^(*IC*_50_*/EC*_50_*/K*_*i*_*/K*_*d*_)). After cleaning and preprocessing (Appendix A.1), we obtained 2,727,579 pairs from 1,018,191 unique compounds and 6,342 unique proteins, split 70/10/20 into train/validation/test sets (random seed 5).

Protein sequences were represented using cached mean embeddings from ESM-C (600M parameters; 1152-d)[6]. Molecules were encoded as DGL graphs with 58-d node features (atomic number, degree, hybridization, etc.) and 13-d edge features (bond type, conjugation, stereochemistry)[11, 12]. Details in Appendix A.2 and A.3.

For screening, we curated CysDB (cysteine-reactive electrophiles) and MolGlueDB (n=1,523 molecular glues)[1, 2].

### 3.2 Model Architecture

The TubercuProbe architecture integrates molecular graphs and protein embeddings through bidirectional cross-attention. The molecular encoder uses five GINEConv layers with learnable *ϵ*, residual connections (layers 2–5), batch normalization, GELU, and dropout [13]. Edge features are projected at each layer to 512-d hidden dimension[14]. Adaptive pooling concatenates mean/max/sum over nodes and edges, projected to 256-d. Protein embeddings (1152-d) are projected to the same 256-d space via a two-layer MLP.

A two-layer bidirectional cross-attention module (16 heads) refines both modalities: self-attention followed by cross-attention (protein↔molecule), with feed-forward blocks (4*×* expansion) and sigmoidal gating for fusion. A three-block MLP head outputs continuous pChEMBL values. Full mathematical details in Appendix A.4. Architecture shown in Figure 1.

**Figure 1.**
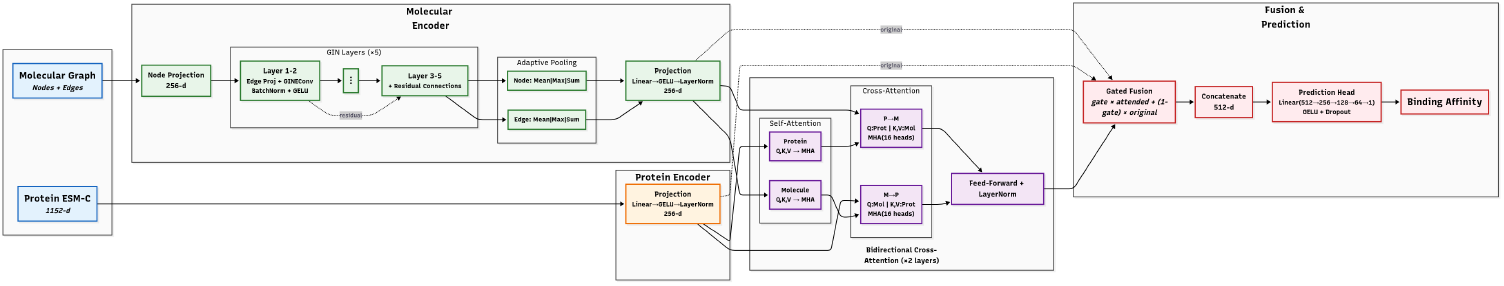
TubercuProbe architecture. Molecular encoder: 5-layer GINE with residual connections and adaptive pooling. Protein encoder: ESM-C mean-pooled embeddings projected to 256-d. Fusion: bidirectional cross-attention with self-attention refinement and sigmoidal gating. Prediction head: 3-block MLP for pChEMBL regression.

### 3.3 Training and Evaluation

Training minimized MSE on pChEMBL values using AdamW (lr=1e-3, weight decay=1e-5) with ReduceLROnPlateau scheduler (patience=20, factor=0.5). Hyperparameters: hidden size 512, batch size 512, dropout 0.23, gradient clipping (max norm 1.0), early stopping (patience=150). Metrics: *R*^2^, MSE, RMSE, median absolute error. Implementation: PyTorch, DGL, RDKit, ClearML. Training on RTX A6000 (48GB) took 24 hours. Full training details in Appendix A.5.

## 4 Results

### 4.1 Overall Regression Performance

On ChEMBL held-out validation and test sets, TubercuProbe achieves *R*^2^ = 0.77, MSE=0.45, indicating strong monotonicity suitable for screening and ranking tasks, surpassing GraphDTA baselines (Table 1). Learning curves depict stable convergence with minimal overfitting.

**Table 1.** Performance comparison of GraphDTA baseline models and TubercuProbe (ours).

| Model | MSE | $R^2$ | Pearson | Median-AE |
| --- | --- | --- | --- | --- |
| GCNNet | 0.81 | 0.59 | 0.77 | — |
| GATNet | 1.12 | 0.42 | 0.65 | — |
| GAT_GCN | 1.09 | 0.44 | 0.66 | — |
| GINConvNet | 1.13 | 0.42 | 0.65 | — |
| **TubercuProbe** | **0.45** | **0.77** | **0.88** | **0.36** |

Both held-out validation and test sets depict approximately normal, unimodal residual histograms (Figure 2), suggesting minimal bias as a predictor. The validation loss curve shows steady convergence, reaching a minimum validation loss of 0.4524 at epoch 679. Occasional spikes in the curve can be attributed to noise in the dataset and the optimizer’s traversal through the non-convex loss landscape.

**Figure 2.**
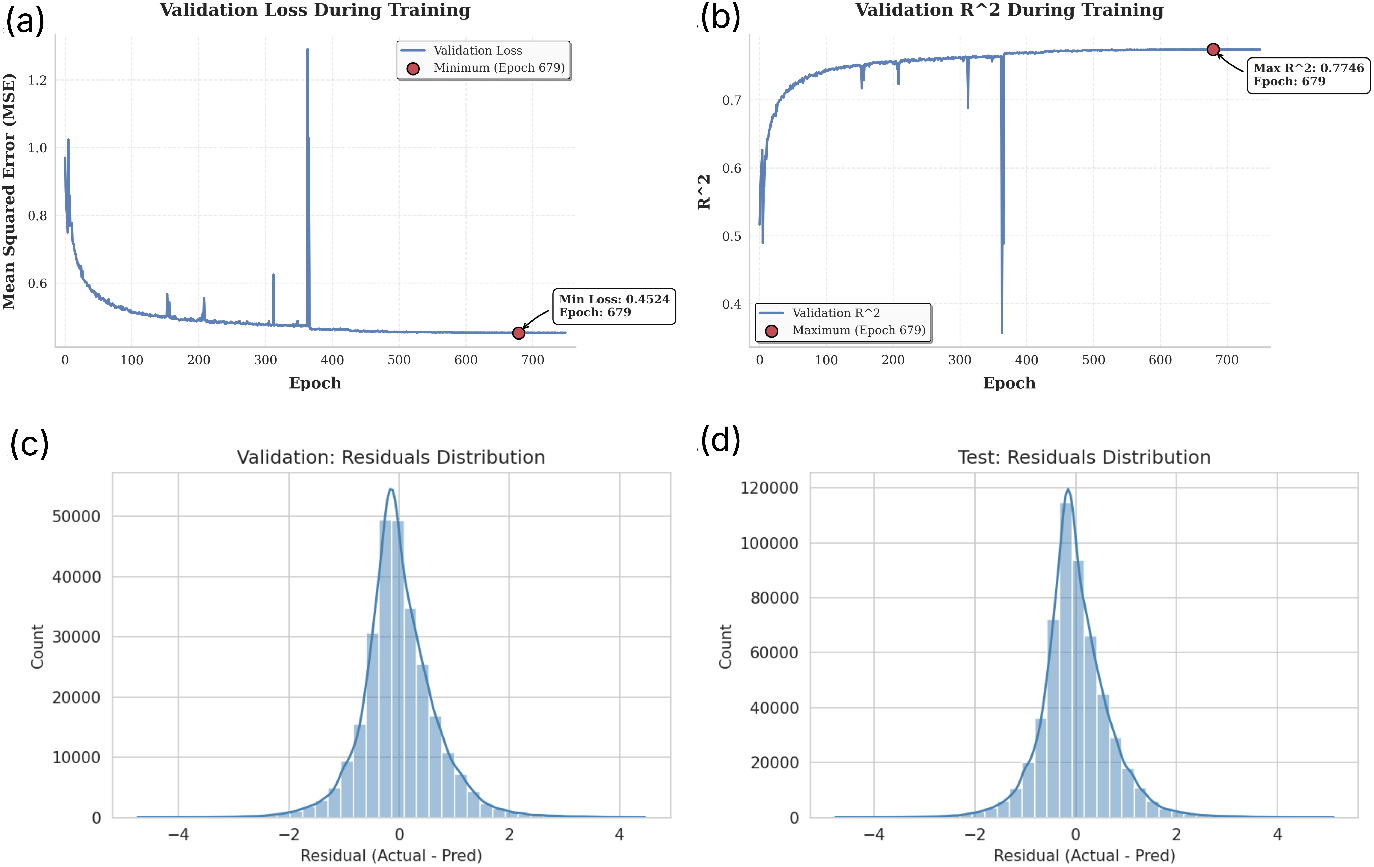
(a) Validation loss curve showing convergence to minimum at epoch 679; (b) Validation *R*^2^ progression; (c) Validation residual histogram; (d) Test residual histogram. Both residual distributions are approximately normal and unimodal, indicating unbiased predictions.

### 4.2 Transfer Learning to CysDB Cysteine Reactivity

Effective covalent probe prioritization requires predicting covalent reactivity, not just non-covalent binding affinity. We fine-tuned TubercuProbe on CysDB [1], a database of cysteine-reactive compounds with binary activity labels from activity-based protein profiling (ABPP) experiments.

#### Dataset

CysDB contains compound-cysteine pairs labeled as reactive (1) or non-reactive (0) based on chemoproteomic measurements. We applied scaffold splitting to ensure no structural overlap between training and test sets, providing rigorous evaluation of generalization to novel chemotypes. Data leakage analysis confirmed 0% similarity between test and training actives (Figure 3).

**Figure 3.**
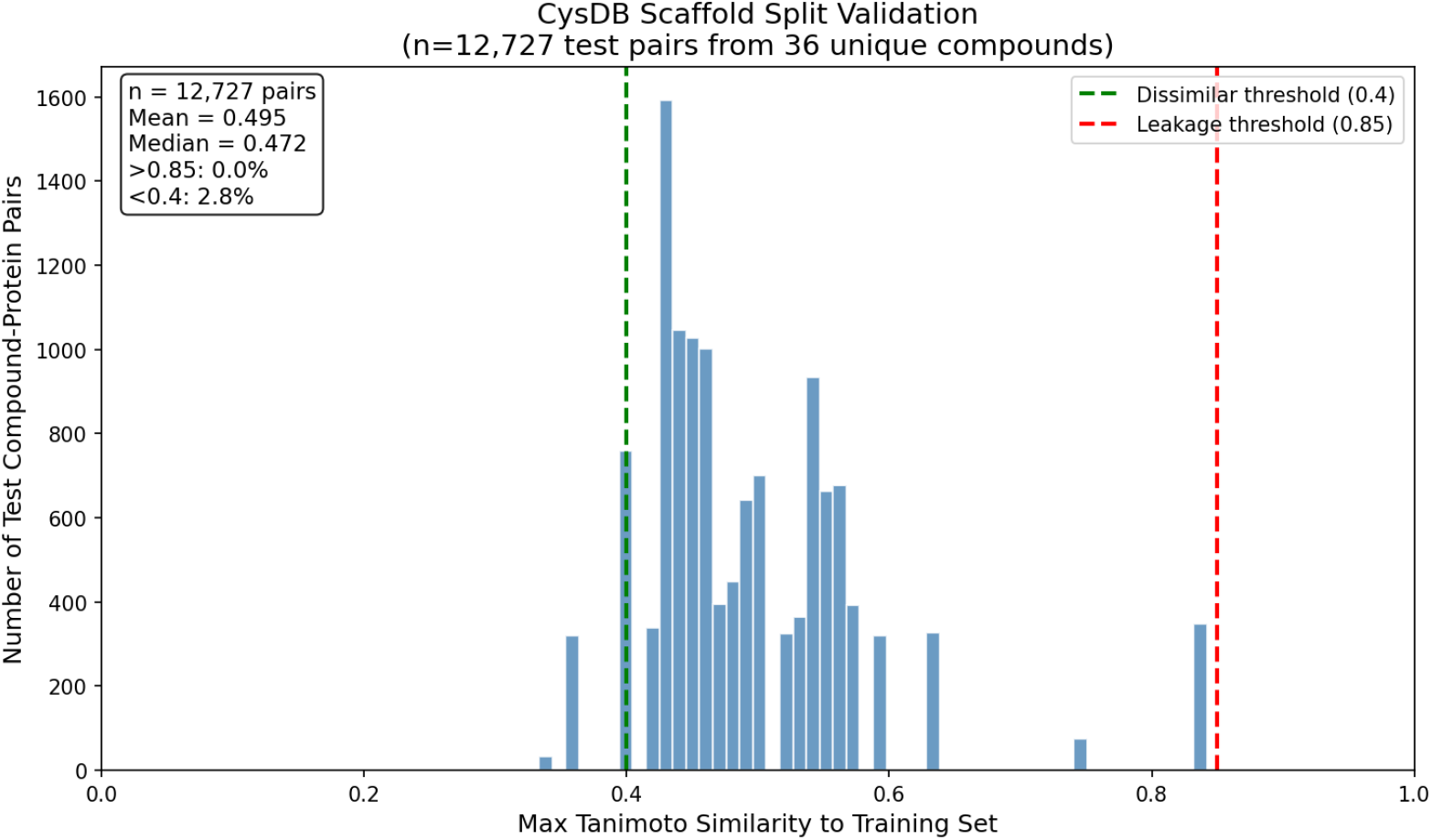
Scaffold split validation for CysDB fine-tuning data (n=12,727 test pairs from 36 unique compounds). Tanimoto similarity distribution between each test compound and its nearest neighbor in the training set. Mean similarity = 0.50, with 0% of pairs exceeding the 0.85 leakage threshold. This confirms that scaffold splitting effectively separates chemotypes, ensuring test performance reflects generalization to novel scaffolds.

#### Fine-tuning protocol

The ChEMBL-pretrained backbone was adapted using binary cross-entropy loss optimized for AUPRC (imbalanced classification). We evaluated eight freezing strategies to understand which model components require task-specific adaptation (Table 2, Figure 6).

**Table 2.** Layer freezing ablation study on CysDB. Strategies are ordered by test AUPRC. Top section: pretrained fine-tuning strategies. Bottom section: controls demonstrating pretraining value.

| Freezing Strategy | Test AUPRC | Test AUROC | Test Acc. | Test F1 |
| --- | --- | --- | --- | --- |
| *ChEMBL-pretrained fine-tuning strategies* |  |  |  |  |
| 1 Freeze cross-attention | **0.6332** | 0.8684 | 0.8031 | 0.5937 |
| 2 Full fine-tune | 0.6327 | 0.8678 | 0.7949 | 0.5862 |
| 3 Freeze GIN early (layers 1–2) | 0.6277 | **0.8698** | 0.8044 | 0.5958 |
| 4 Freeze input projections | 0.6270 | 0.8680 | 0.7996 | 0.5889 |
| 5 Freeze all GIN (layers 1–5) | 0.6078 | 0.8657 | **0.8053** | 0.5926 |
| *Controls (limited adaptation capacity)* |  |  |  |  |
| 6 Adapt attention only (freeze GIN + proj.) | 0.4519* | — | — | — |
| 7 Adapt late GIN + attention | 0.4518* | — | — | — |
| 8 Feature extraction (train head only) | 0.4502* | — | — | — |

\*Validation AUPRC reported (experiments did not complete to test evaluation).

**Table 3.** Complete hyperparameter configuration.

| Hyperparameter | Value |
| --- | --- |
| Node feature dim | 58 |
| Edge feature dim | 13 |
| GINE hidden dim | 512 |
| Number of GINE layers | 5 |
| Protein embedding dim | 1152 |
| Latent embedding dim | 256 |
| Cross-attention layers | 2 |
| Attention heads | 16 |
| Batch size | 512 |
| Learning rate | $1 \times 10^{-3}$ |
| Weight decay | $1 \times 10^{-5}$ |
| Dropout rate | 0.23 |
| Gradient clip norm | 1.0 |
| Early stopping patience | 150 |
| LR scheduler patience | 20 |
| Max epochs | 3000 |

#### Key findings

The ablation study reveals two important patterns:

1. **High transferability of pretrained features**: The narrow performance range (ΔAUPRC=0.025) across fine-tuning strategies (rows 1–5) demonstrates that ChEMBL-pretrained molecular representations generalize effectively to covalent reactivity prediction. Freezing cross-attention achieves marginally the *best* performance, suggesting that protein-molecule interaction patterns learned from non-covalent binding transfer directly.
2. **Pretraining provides substantial benefit**: The control experiments (rows 6–8) show dramatic performance drops when adaptation capacity is severely limited. Feature extraction (train head only) achieves only AUPRC=0.45, compared to 0.63 for fine-tuning strategies—a 40% relative improvement. This gap confirms that pretrained backbone features, not just the classification head, are critical for performance.

A visualization of which layers are frozen in each strategy is provided in Appendix A.10.

#### Implications for multitask learning

These results motivate a multitask extension where the model jointly predicts non-covalent affinity and covalent reactivity. Effective covalent probes must both *reach* their target (binding affinity) and *react* once there (covalent reactivity). Preliminary data preparation combining ChEMBL affinity data with CysDB reactivity labels is complete; multitask training is an active area of development.

### 4.3 Screening Compound Libraries: CysDB and Molecular Glues

Mycobacterium tuberculosis is the world’s leading bacterial killer, with mortality projected to rise as resistance spreads. Its secreted effector proteins (PtpB, Rv3671c, SapM) are central to immune evasion and represent underexplored targets.

Activity-based protein profiling (ABPP) maps protein-probe interactions using covalent probes, but campaigns are costly and low-throughput. TubercuProbe enables large-scale *in silico* screening of molecular probes, prioritizing candidates for experimental follow-up. We screened CysDB (cysteine-reactive warheads) and MolGlueDB against three Mtb targets, reporting top-ranked compounds. Screening protocols detailed in Appendix A.8.

### 4.4 Comparison with Boltz-2

Our results show moderate correlation between TubercuProbe and Boltz-2 predictions. TubercuProbe outputs pChEMBL values (− log_10_(affinity), higher = stronger binding), while Boltz-2 outputs log affinity (higher = weaker binding). After accounting for this sign convention, the combined correlation across CysDB and MolGlueDB datasets (|*r*| ≈ 0.69, *p <* 10^−9^; Figure 4) indicates substantial overall agreement between the sequence-graph and structure-based approaches. *Note: this correlation is computed across combined datasets; per-dataset correlations may differ*. Boltz-2 requires minutes per inference; our model achieves sub-second predictions, enabling proteome-wide screening. TubercuProbe handles intrinsically disordered proteins where structural methods struggle. While Boltz-2 reports *R*^2^ = 0.69 on FEP+ benchmarks[7], our approach offers cost-efficient, sequence-driven ranking at scale, with structure-based models reserved for high-resolution follow-up.

**Figure 4.**
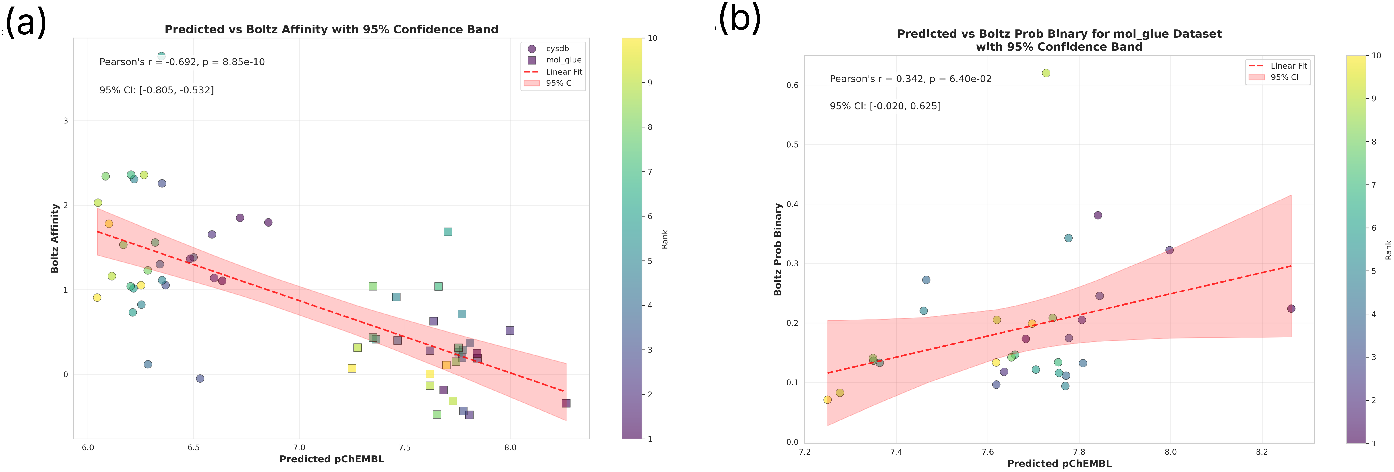
(a) Predicted affinity scatter comparing TubercuProbe vs. Boltz-2 on top-10 compounds per target (CysDB: circles, MolGlueDB: squares). Combined Pearson *r* = −0.69 reflects opposite sign conventions between models. (b) Correlation between affinity classes showing consistent agreement.

## 5 Discussion and Limitations

### Why regression helps ranking

Pointwise pChEMBL regression yields calibrated, continuous scores trivially sortable across proteins, enabling simple top-k triage. It also supports gradient analyses and fragment-level rationales via substructure-masking[15].

### Limitations

(i) Training on ChEMBL may underperform on extreme chemotypes; (ii) no explicit 3D structure; (iii) no calibrated uncertainty; (iv) fine-tuning on chemoproteomic data remains challenging due to binary/ratio readouts.

### Positioning

Boltz-2 achieves near-FEP correlations with complex generation[8] but at higher cost. TubercuProbe offers cheaper, high-throughput ranking complementary to downstream co-folding—analogous to SimpleFold’s efficient approach[9].

### 6 Broader Impact and Ethics

Sequence-based screening can accelerate probe discovery and repurposing. Potential negative uses include dual-use for pathogen research; we release code with usage guidelines and recommend controlled access. We do not model off-targets, toxicity, or ADME; experimental validation is required. Prioritization of covalent probes and molecular glues could accelerate anti-tubercular target validation, reducing computational, economic, and time costs. TubercuProbe operationalizes a graph-sequence cross-attention pipeline for proteome-scale screening with interpretable attention and full reproducibility.

#### Note

This preprint reflects ongoing work. *In silico* experiments and prospective ABPP validation are in progress; this paper will be updated incrementally as new results become available.

## A Technical Appendix

This appendix provides comprehensive technical details for exact replication, including data processing, feature engineering, architectural specifications, training procedures, and implementation notes. The github repository contains all scripts and notebooks to reproduce the results. https://github.com/ABHCI15/AffinityPred.

### A.1 Data Curation and Preprocessing

#### Source and filtering

ChEMBL database[10] version 33 was queried for compound-protein affinity measurements with standardized pChEMBL values. We filtered for assays with defined *IC*_50_, *EC*_50_, *K*_*i*_, or *K*_*d*_ measurements and converted to pChEMBL scale: pChEMBL = − log^10^(affinity in M).

#### Molecule standardization

(i) SMILES strings were sanitized using RDKit’s `Chem.MolFromSmiles()` with error handling; (ii) salts and counterions were removed using `rdMolStandardize.LargestFragmentChooser();` (iii) invalid molecules (failed sanitization, molecular weight *>*1000 Da, or containing rare elements) were discarded; (iv) remaining molecules were canonicalized using `Chem.MolToSmiles(canonical=True)`.

#### Protein mapping

Protein targets were mapped to UniProt accessions via ChEMBL target dictionary. Sequences were retrieved from UniProt with minimum length 50 and maximum length 2000 residues. Proteins with missing or ambiguous sequences were excluded.

#### Label processing

pChEMBL values were clipped to [1st percentile, 99th percentile] = [4.12, 10.87] to reduce the impact of outlier measurements and potential annotation errors. This resulted in a total of 2,727,579 unique compound-protein pairs across 1,018,191 unique compounds and 6,342 unique proteins.

#### Data splits

Random stratified split with 70% training (1,909,305 pairs), 10% validation (272,758 pairs), and 20% test (545,516 pairs), using random seed 5 for reproducibility. No protein or compound leakage between splits was enforced at the pair level.

#### Screening datasets

(i) CysDB: 300 cysteine-reactive electrophiles with annotations for ligandability scores and hyperreactivity flags; (ii) MolGlueDB: 1,523 molecular glue degraders with E3 ligase and target protein annotations. Both datasets were processed using identical molecule standardization pipelines.

### A.2 Molecular Graph Features

#### Node features (58-dimensional)

Each atom is represented by a concatenated feature vector:

- Atomic number (1-d, integer)
- Degree (1-d, total number of bonds)
- Total hydrogen count (1-d, explicit + implicit)
- Formal charge (1-d, integer)
- Aromaticity (1-d, binary)
- Ring membership (1-d, binary)
- Atom type one-hot (44-d, covering common organic elements + “Unknown”)
- Hybridization one-hot (7-d: SP, SP2, SP3, SP3D, SP3D2, S, Other)
- Number of radical electrons (1-d)

#### Edge features (13-dimensional)

Each bond is represented by:

- Bond type one-hot (4-d: single, double, triple, aromatic)
- Conjugated (1-d, binary)
- In ring (1-d, binary)
- Stereo flag (1-d, binary)
- Stereo configuration one-hot (6-d: none, any, Z, E, cis, trans)

#### Graph construction

Molecular graphs are constructed using DGL with bidirectional edges. For a molecule with *N* atoms and *M* bonds, the graph *G* = (*V, E*) has |*V* | = *N* nodes and |*E*| = 2*M* directed edges (each undirected bond becomes two directed edges). Node features are stored as `g.ndata[‘feat’] and edge features as g.edata[‘feat’]`.

#### Implementation details

Graph construction uses RDKit’s `GetAtoms()` and `GetBonds()` with feature extraction via custom feature functions. Graphs are batched using DGL’s `dgl.batch()` for efficient GPU processing.

### A.3 Protein Sequence Embeddings

#### ESM Cambrian (ESM-C) model

We use ESM-C 600M, part of the ESM Cambrian family released by EvolutionaryScale in December 2024 [16]. Unlike its predecessor ESM2, ESM-C is trained on a substantially larger and more diverse dataset comprising UniRef, MGnify, and Joint Genome Institute (JGI) metagenomic sequences, totaling over 2.4 billion clustered sequences at 70% sequence identity. The 600M parameter model uses a transformer architecture with 36 layers, width 1152, and 18 attention heads, employing Pre-LN, rotary positional embeddings, and SwiGLU activations. The model produces per-residue embeddings of dimension 1152.

#### Embedding generation

(i) Protein sequences are tokenized using ESM-C’s built-in tokenizer; (ii) sequences are passed through the model to obtain per-residue representations **H** ∈ ℝ^*L×*1152^ where *L* is sequence length; (iii) mean pooling is applied across the sequence dimension to obtain a fixed 1152-d vector: 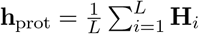.

#### Caching strategy

All protein embeddings are precomputed offline using the AffinityDataProcessor class and cached as pickled dataframes to avoid redundant computation during training. The processor initializes ESM-C lazily and maintains an embedding cache that is saved periodically.

#### Handling long sequences

For proteins exceeding ESM-C’s maximum context length (2048 tokens), we use sliding window averaging with 50% overlap and aggregate via mean pooling.

### A.4 Detailed Architecture Specifications

#### A.4.1 Graph Neural Network Background

##### From QSAR to GNNs

Traditional Quantitative Structure-Activity Relationship (QSAR) models utilize mathematical relationships between chemical structures and biological activities based on handcrafted molecular descriptors (molecular weight, LogP, topological indices). While interpretable and using techniques like multiple linear regression (MLR) and partial least squares (PLS), QSAR assumes linear relationships and requires computation of thousands of descriptors that may fail to capture intricate nonlinear patterns in drug-target interactions.

Graph Neural Networks (GNNs) overcome these limitations by learning features directly from graph-structured molecular data[17]. Molecules are naturally represented as graphs with atoms as nodes and bonds as edges. GNNs iteratively update node embeddings based on local graph structure, capturing complex hierarchical relationships. [13]

##### General GNN aggregation. The standard GNN update rule is

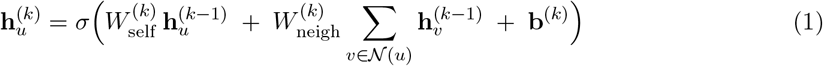

where 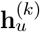 is the node *u* embedding at layer *k, σ* is a nonlinearity, 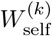 and 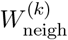 are learnable weights, 𝒩 (*u*) denotes neighbors of *u*, and **b**^(*k*)^ is bias.

#### A.4.2 Why GINE over GCN, GAT, and GIN Limitations of GCN and GAT

- **GCN**: Suffers from oversmoothing with deep layers, uses fixed aggregation weights (all neighbors contribute equally), not injective (different structures → same embeddings), poor scalability to large graphs.
- **GAT**: Improves via learned attention weights but adds computational overhead, still not maximally expressive, lacks injectivity guarantees.

##### GIN advantages

Graph Isomorphism Networks achieve maximal expressiveness by ensuring injective aggregation functions, matching the discriminative power of the Weisfeiler-Lehman (WL) graph isomorphism test[14]. The GIN update rule is:

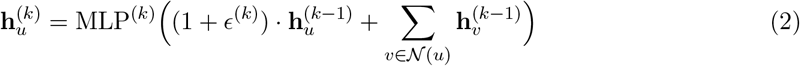

where *ϵ*^(*k*)^ is a learnable scalar weighting the central node. The sum aggregation plus learnable *ϵ* and MLP guarantee injectivity, enabling GIN to distinguish subtle structural differences (e.g., isomers) critical in drug discovery.

##### GINE extension

Standard GIN ignores edge features, limiting expressiveness for molecules where bond types (single/double/aromatic) are essential. GINE (Graph Isomorphism Network with Edge features) extends GIN by incorporating edge-aware aggregation:

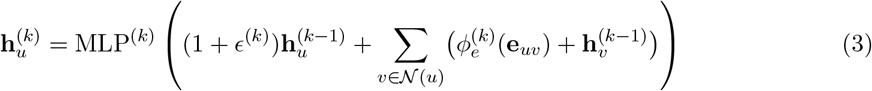

where 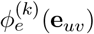 projects edge features into the node feature space. This allows GINE to leverage bond-level information (conjugation, stereochemistry) alongside atomic properties, yielding superior performance on molecular property prediction tasks.

#### A.4.3 Molecular Encoder: GINE Layers with Residual Connections

Our architecture employs five GINE layers with architectural enhancements for stability and expressiveness.

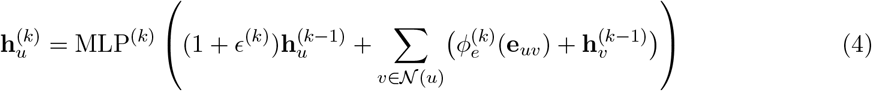

##### GINE update rule. For layer *k* and node *u*

##### Layer-specific components

Each GINE layer *k* includes:

- **Edge projection**: 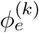 : ℝ^13^ → ℝ^512^ (linear layer mapping 13-d bond features to hidden dimension)
- **Node MLP**: Two-layer perceptron with 512-d hidden dimension, batch normalization after each linear layer, GELU activation, and dropout (0.23):

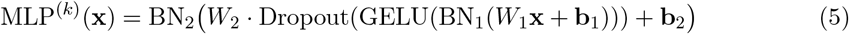
- **Learnable** *ϵ*^(*k*)^: Initialized and optimized per layer, allowing adaptive weighting of self vs. neighbor information
- **Batch normalization**: Applied after GINE aggregation to stabilize training
- **Residual connections (layers 2–5)**: **h**^(*k*)^ ← **h**^(*k*)^ + **h**^(*k*−1)^ to mitigate gradient vanishing and enable deeper architectures

#### Rationale for residual connections

Deep GNNs can suffer from oversmoothing (node embeddings become indistinguishable). Residual connections preserve information from earlier layers, allowing the network to learn hierarchical representations at multiple scales. Layers 2–5 include residuals; layer 1 establishes initial embeddings without prior context.

##### Initial projection

Input node features (58-d) are projected to 512-d via:

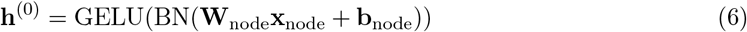

##### Forward pass through GINE stack

The complete forward pass is:

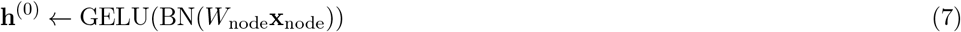

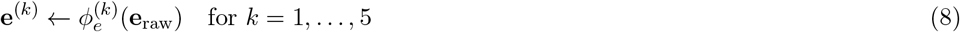

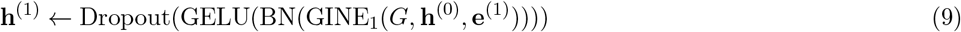

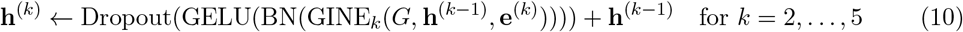

##### Computational complexity

Each GINE layer requires *O*(|*E*| · *d*^2^) operations for edge projection and MLP application, where |*E*| is the number of edges and *d* = 512 is hidden dimension. Total complexity for five layers is *O*(5|*E*|*d*^2^), manageable for molecular graphs (typically |*E*| *<* 100).

#### A.4.4 Adaptive Graph Pooling

After the final GINE layer, we obtain node representations 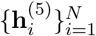 and edge representations.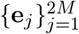. Adaptive pooling aggregates these into fixed-size graph embeddings.

##### Motivation

Molecular graphs vary in size (small fragments vs. large macrocycles). Graph-level predictions require fixed-size representations. Standard pooling (mean/max/sum) captures different aspects: mean preserves average properties, max highlights salient features, sum reflects graph size. Combining all three yields richer representations.

##### Node pooling

Three pooling operations are applied:

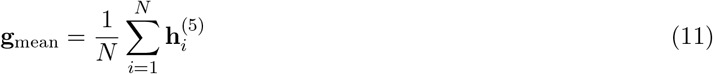

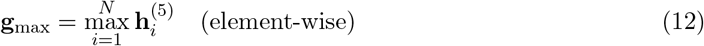

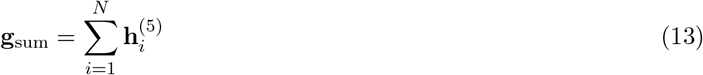

##### Edge pooling

Similarly for edges (capturing bond-level information):

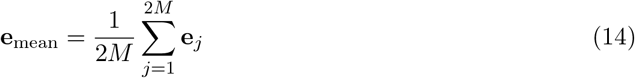

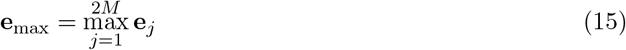

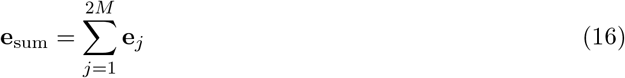

##### Combined representation

Node and edge pools are concatenated and projected:

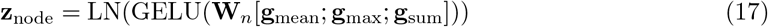

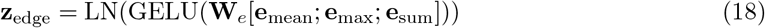

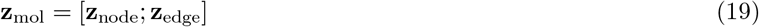

where **W**_*n*_ : ℝ^1536^ → ℝ^256^ (3*×*512-d node pools to 256-d), **W**_*e*_ : ℝ^39^ → ℝ^256^ (3*×*13-d edge pools to 256-d), and LN denotes LayerNorm. The final concatenation yields a 512-d molecular embedding.

##### Final molecular projection

The concatenated 512-d representation is projected to 256-d:

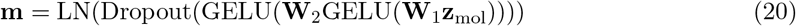

where **W**_1_ : ℝ^512^ → ℝ^512^ and **W**_2_ : ℝ^512^ → ℝ^256^.

#### A.4.5 Protein Encoder

Protein sequences are encoded using ESM-C 600M [16], a next-generation protein language model from the ESM Cambrian family (EvolutionaryScale, December 2024). Unlike ESM2, ESM-C is trained on a massively expanded dataset including UniRef (83M clusters), MGnify (372M clusters), and JGI metagenomic sequences (2B clusters). The model is trained using masked language modeling on 6.2 trillion tokens over 1.5M steps. ESM-C leverages a modern transformer architecture with Pre-Layer Normalization, rotary positional embeddings, and SwiGLU activations, achieving significant performance improvements over ESM2 with equivalent or reduced computational cost.

##### Why ESM-C?

Protein binding sites exhibit complex physicochemical patterns (hydrophobic patches, electrostatic complementarity, hydrogen bond donors/acceptors) not easily captured by sequence alone. ESM-C embeddings encode: (1) evolutionary conservation (functional residues), (2) co-evolution patterns (structural contacts), (3) local structural preferences (secondary structure propensities), and (4) long-range dependencies (domain interactions). The expanded metagenomic training data provides richer representations for uncommon protein families, including pathogen proteins with limited homologs in UniRef.

##### Embedding extraction

For a protein sequence *S* = (*s*_1_, …, *s*_*L*_) of length *L*, ESM-C produces per-residue embeddings 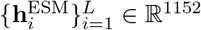. We apply mean pooling over the sequence:

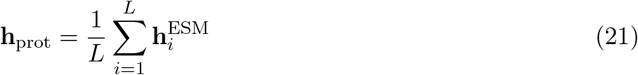

##### Projection and normalization

The protein encoder projects the pooled 1152-d ESM-C embedding to 256-d through a 2-layer MLP with intermediate expansion to 512-d:

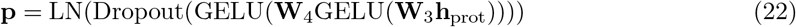

where **W**_3_ : ℝ^1152^ → ℝ^512^ and **W**_4_ : ℝ^512^ → ℝ^256^. LayerNorm (LN) stabilizes training, Dropout (0.23) prevents overfitting to protein families, and GELU provides smooth non-linearity.

##### Computational efficiency

ESM-C 600M achieves high throughput (∼40-70 sequences/second on GPU), making real-time inference feasible. However, we pre-compute and cache all protein embeddings during dataset preparation (see Appendix A.1) to avoid redundant computation across training epochs, reducing per-epoch time from minutes to ∼30ms per batch.

#### A.4.6 Bidirectional Cross-Attention

The cross-attention module enables bidirectional information exchange between molecular and protein representations. This fusion mechanism allows the model to learn complementary binding patterns: molecules query protein binding sites for pharmacophore compatibility, while proteins query molecular substructures for functional group interactions.

##### Architecture overview

The module consists of two stacked layers (*L* = 2), each with 16 attention heads (*H* = 16, head dimension *d*_*k*_ = 16). Each layer performs: (1) intra-modality self-attention to refine individual representations, (2) inter-modality cross-attention for bidirectional fusion, and (3) feed-forward networks for non-linear transformation.

##### Layer 1: Self-attention

Both modalities first undergo self-attention to capture intramodality dependencies (e.g., correlations between different molecular subgraphs, or protein domain interactions):

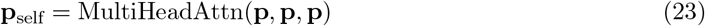

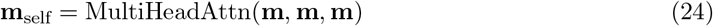

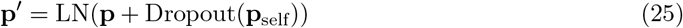

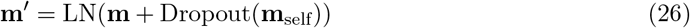

##### Layer 2: Cross-attention

Protein queries molecule (what molecular features are relevant for this protein?) and molecule queries protein (what protein regions are compatible with this ligand?):

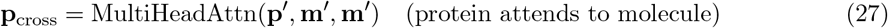

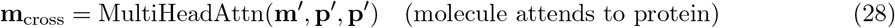

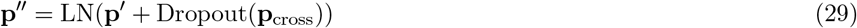

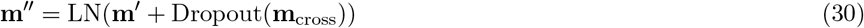

##### Layer 3: Feed-forward networks

Per-modality FFNs with 4*×* expansion (256 → 1024 → 256) add non-linear expressiveness:

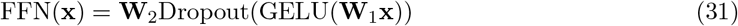

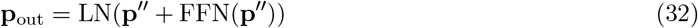

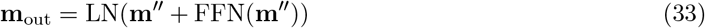

where **W**_1_ : ℝ^256^ → ℝ^1024^ and **W**_2_ : ℝ^1024^ → ℝ^256^.

##### Multi-head attention mechanism

For query **Q**, key **K**, value **V**, the multi-head attention with *H* = 16 heads computes:

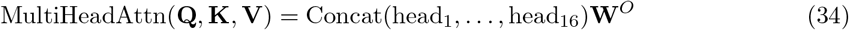

where each head *h* independently computes scaled dot-product attention:

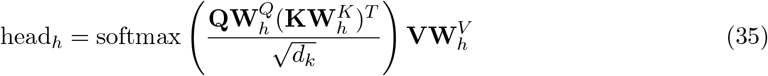

with head dimension *d*_*k*_ = 256*/*16 = 16. Projection matrices 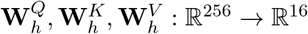 project to head-specific subspaces, and **W**^*O*^ : ℝ^256^ → ℝ^256^ combines heads.

##### Why bidirectional?

Unidirectional attention (only molecule→protein or protein→molecule) misses symmetrical binding complementarity. Bidirectional cross-attention captures: (1) molecular features relevant to protein pockets (e.g., hydrophobic groups aligning with lipophilic residues), (2) protein features relevant to ligands (e.g., charged residues attracting polar groups), and (3) mutual structural complementarity (shape and electrostatic fit). This symmetry improves generalization to unseen protein-ligand pairs [5].

#### A.4.7 Gated Fusion

After the cross-attention layers, we fuse the original embeddings (**p**_orig_, **m**_orig_) with the attended embeddings (**p**_out_, **m**_out_) via a learnable gating mechanism. This allows the model to adaptively balance original representations (preserving intrinsic molecular/protein properties) with cross-attended representations (encoding binding complementarity).

##### Gating mechanism

A sigmoid-activated gate **g** ∈ [0, 1]^256^ controls the fusion ratio:

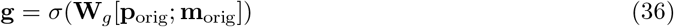

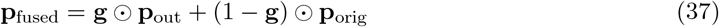

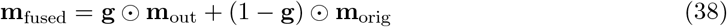

where *σ* is the sigmoid function, **W**_*g*_ : ℝ^512^ → ℝ^256^ projects concatenated original embeddings, and ⊙ denotes element-wise multiplication. When **g** ≈ 1, the model trusts cross-attention (strong protein-ligand interaction); when **g** ≈ 0, it preserves original features (weak or no interaction).

##### Why gated fusion?

Simple concatenation or addition treats all features equally. Gating enables the model to: (1) downweight noisy cross-attention signals for dissimilar protein-ligand pairs, (2) amplify relevant cross-modal interactions for binding pairs, and (3) learn feature-specific fusion strategies (e.g., preserve hydrophobicity from molecule, emphasize charged residues from protein).

#### A.4.8 Prediction Head

The prediction head maps the concatenated fused embeddings (512-d) to a scalar pChEMBL prediction via a 3-layer MLP with progressive dimensionality reduction.

##### Architecture

Three hidden layers with exponentially decreasing width (512 → 256 → 128 → 64):

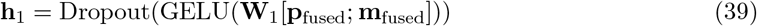

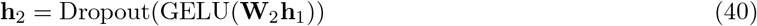

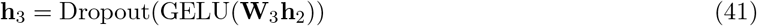

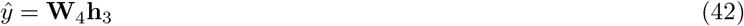

where **W**_1_ : ℝ^512^ → ℝ^256^, **W**_2_ : ℝ^256^ → ℝ^128^, **W**_3_ : ℝ^128^ → ℝ^64^, **W**_4_ : ℝ^64^ → ℝ^1^ (linear output, no activation).

##### Design rationale

The funnel architecture progressively distills joint protein-ligand features into a scalar affinity score. Dropout (0.23) at each layer prevents co-adaptation of hidden units. GELU activations provide smooth gradients. The linear final layer enables unconstrained regression (pChEMBL ranges 4–12, requiring unbounded outputs).

##### Total parameters

The entire model contains ∼6.9M trainable parameters: molecular encoder, protein encoder, cross-attention, fusion gate, prediction head. This compact size enables training ona single GPU while maintaining strong expressiveness.

### A.5 Training Protocol

#### A.5.1 Loss Function and Optimization

**Objective**. Mean squared error on pChEMBL values:

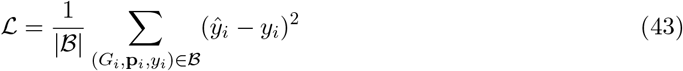

**Optimizer**. AdamW with parameters:

- Learning rate: *α* = 1 *×* 10^−3^
- Weight decay: *λ* = 1 *×* 10^−5^
- Betas: *β*_1_ = 0.9, *β*_2_ = 0.999
- Epsilon: *ϵ* = 1 *×* 10^−8^

**Learning rate schedule**. ReduceLROnPlateau monitors validation RMSE:

- Factor: 0.5 (halve learning rate)
- Patience: 20 epochs
- Cooldown: 10 epochs
- Minimum learning rate: 1 *×* 10^−6^

**Gradient clipping**. Global norm clipping with max norm 1.0 to prevent exploding gradients.

**Early stopping**. Training halts if validation RMSE does not improve for 150 consecutive epochs. Best model checkpoint (lowest validation RMSE) is saved and used for final evaluation.

#### A.5.2 Regularization

- Dropout: 0.23 in all layers
- Feature dropout: 0.115 (50% of base dropout) applied before fusion
- Batch normalization in GINE layers
- Layer normalization in attention and projection modules
- Weight decay in optimizer

#### A.5.3 Hyperparameters Summary

### A.6 Evaluation Metrics

**Mean Squared Error (MSE):**

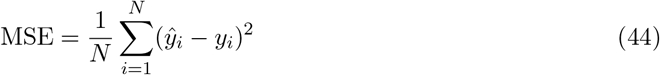

**Root Mean Squared Error (RMSE):**

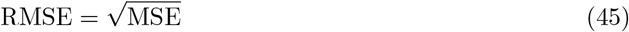

**Coefficient of Determination (***R*^2^):

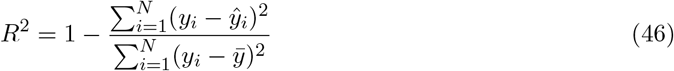

**Pearson Correlation Coefficient:**

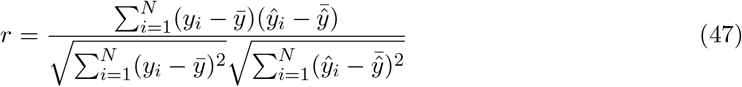

**Median Absolute Error:**

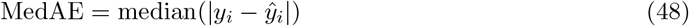

### A.7 Implementation Details

#### Software dependencies

- Python
- PyTorch + CUDA
- DGL
- RDKit
- NumPy
- scikit-learn
- ClearML
- pandas
- matplotlib
- rest are listed in repository

#### Code organization

- `AffinityPred/models/ginconvBiDirectionalNoAttPool.py`: Model architecture
- `AffinityPred/train.py`: Training script
- `AffinityPred/data_processing_utils/`: Data preprocessing utilities
- `AffinityPred/datasets/`: Processed datasets and embeddings
- `AffinityPred/weights/`: Model checkpoints

#### Reproducibility

Random seeds are set for Python (5), NumPy (5), PyTorch (5), and DGL (5). Note that full determinism on GPU is challenging due to CUDA’s non-deterministic operations; results may vary slightly across runs.

#### Data availability

Processed datasets (`train_processed.pkl, valid_processed.pkl, test_processed.pkl`) include precomputed graphs and protein embeddings; if not present, generate with data processing utilities provided. Raw ChEMBL data can be obtained from https://www.ebi.ac.uk/chembl/.

### A.8 Screening Pipeline Details

#### CysDB screening workflow

1. Load 300 cysteine-reactive electrophiles with SMILES and annotations
2. Generate molecular graphs using standard featurization
3. For each target protein (PtpB, Rv3671c, SapM):
  - Load precomputed ESM-C embedding
  - Batch predict pChEMBL scores for all compounds
  - Rank compounds by predicted affinity (descending)
  - Filter by ligandability score *>* 0.5 if applicable
  - Select top-k (k=10, 50, 100) for ABPP follow-up

#### MolGlueDB screening workflow

1. Load 1,523 molecular glues with SMILES and E3 ligase annotations
2. Generate molecular graphs
3. For each target protein:

- Predict affinity for all molecular glues
- Rank by predicted pChEMBL
- Cross-reference with known E3 ligase compatibilities
- Prioritize compounds with favorable ternary complex formation potential

#### Boltz-2 comparison protocol

1. Select top-10 compounds per target from TubercuProbe predictions
2. Prepare inputs: protein FASTA sequence and compound SMILES
3. Run Boltz-2 co-folding (generates 3D structures and predicts affinity internally)
4. Extract predicted affinity from Boltz-2 output
5. Compute Pearson correlation and rank agreement metrics

### A.9 Data Collection Pipeline

Figure 5 illustrates the complete data collection and processing pipeline used in this study.

**Figure 5.**
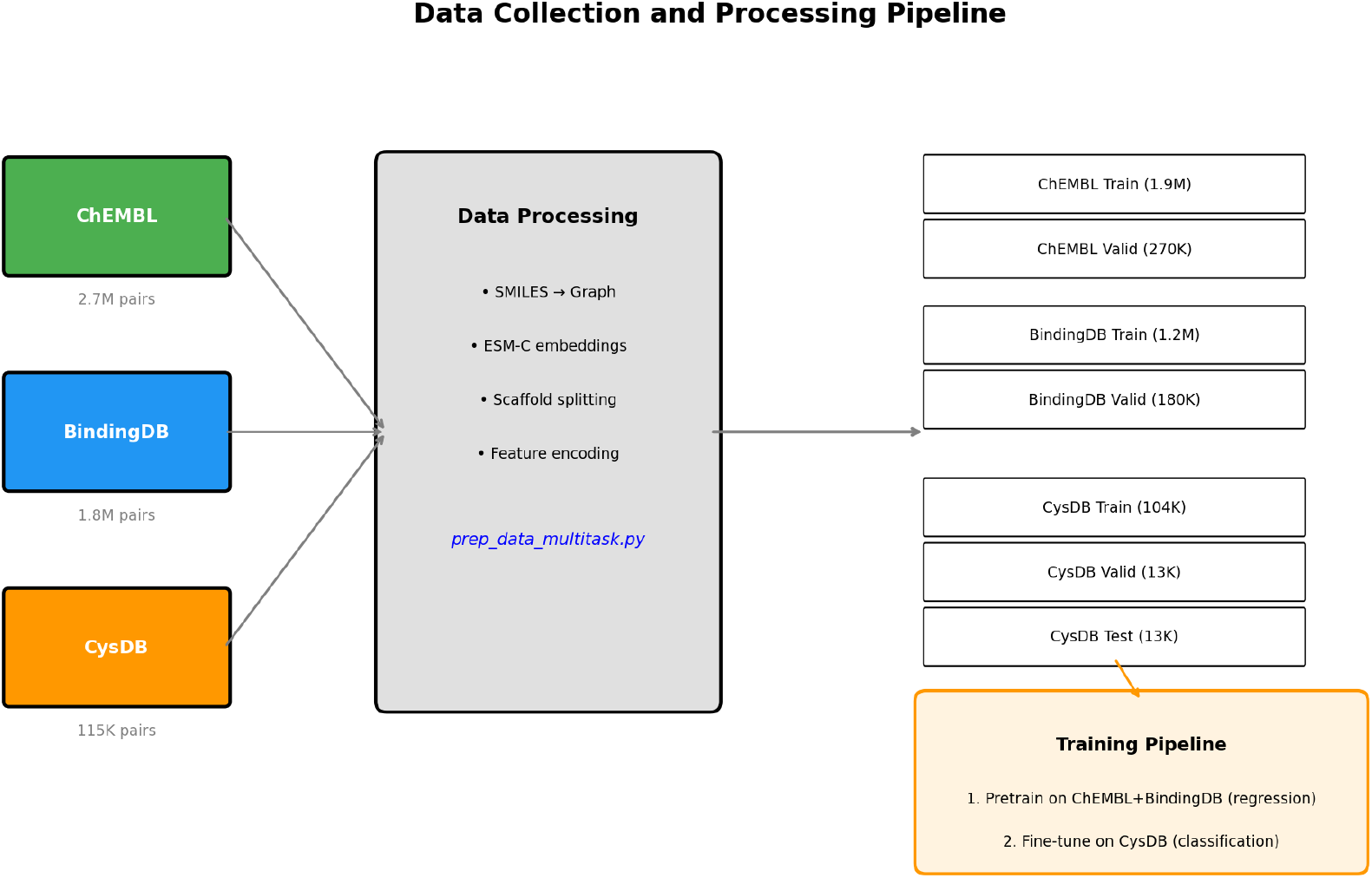
Data collection and processing pipeline. Raw data from ChEMBL (2.7M compound-protein pairs), BindingDB (1.8M pairs), and CysDB (115K pairs) are processed through a unified pipeline that generates molecular graphs, computes ESM-C embeddings, and applies scaffold splitting for rigorous train/test separation.

### A.10 Layer Freezing Ablation Details

Figure 6 shows the decision tree for our layer freezing ablation study, illustrating which components are frozen vs. trainable in each experimental condition.

**Figure 6.**
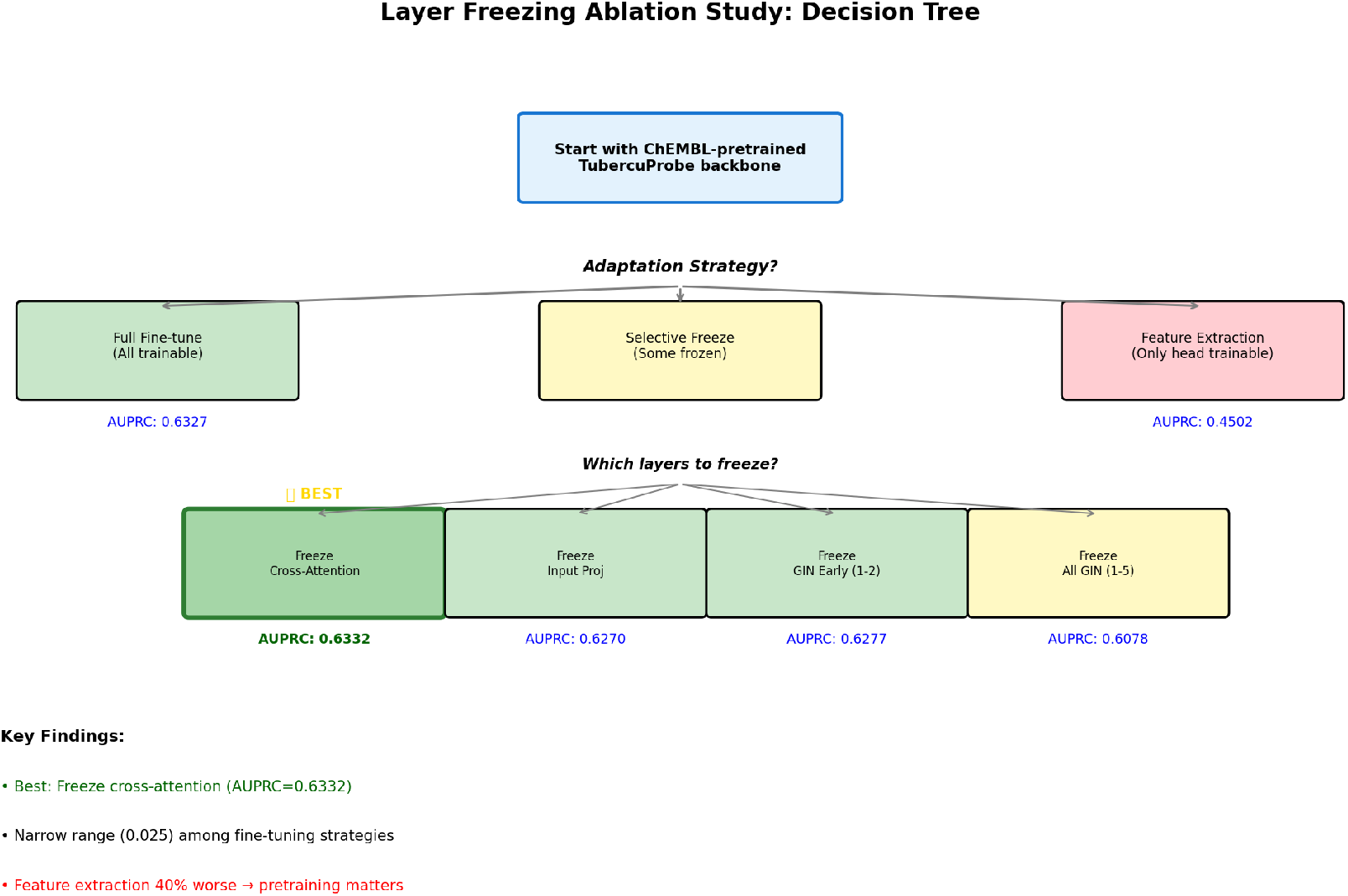
Layer freezing ablation decision tree. Starting from ChEMBL-pretrained weights, we evaluated three adaptation strategies: full fine-tuning (all layers trainable), selective freezing (some layers frozen), and feature extraction (only prediction head trainable). Selective freezing strategies explored which architectural components—GIN layers, input projections, or cross-attention—could be frozen without performance degradation. Key finding: freezing cross-attention achieved the best AUPRC (0.6332), while feature extraction performed 40% worse (0.4502), demonstrating the value of pretrained backbone features.

**Figure 7.**
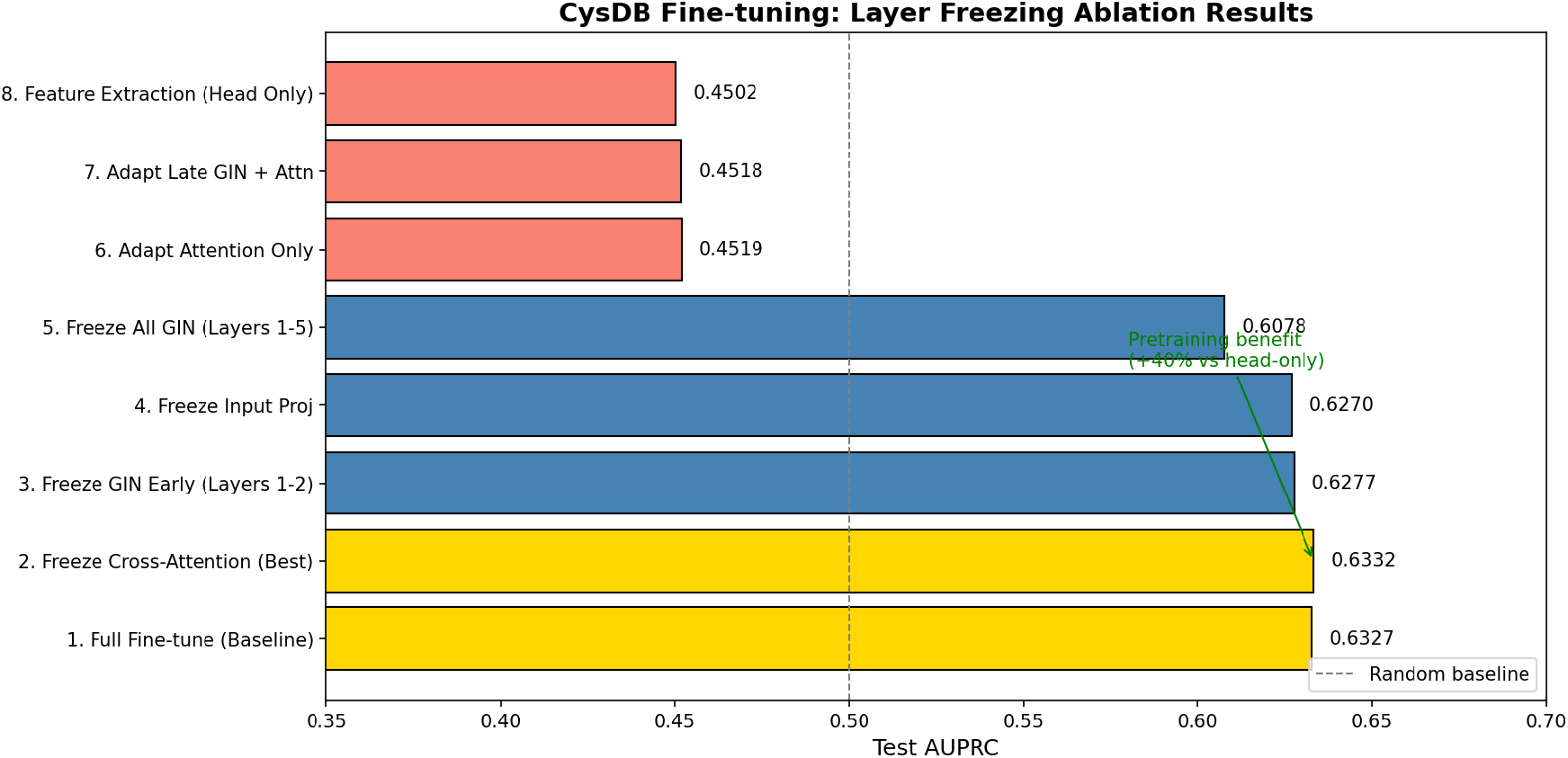
Ablation study results comparison. Horizontal bar chart showing Test AUPRC for all eight freezing strategies. The top five strategies (full fine-tune through freeze all GIN) show a narrow performance range (ΔAUPRC=0.025), indicating high transferability of pretrained features. The bottom three (adapt attention only, adapt late GIN + attention, feature extraction) show substantial drops, confirming that backbone adaptation—not just head training—is critical for CysDB transfer learning.

### A.11 Computational Resources

#### Training infrastructure

- Node: Single HPC compute node
- CPU: 15 cores (AMD Threadripper)
- GPU: 1*×* NVIDIA RTX A6000 (48GB VRAM, Ampere architecture)
- RAM: 120GB DDR4
- Storage: 2TB NVMe SSD for fast data loading

#### Training performance

- Time per epoch: ∼115 seconds (3,730 batches)
- Total training time: ∼24 hours (679 epochs to best model)
- GPU utilization: 85-95%

#### Inference performance

- Single compound-protein pair: ∼2-200 ms (GPU)
- Proteome-scale screening (10K compounds *×* 100 proteins): ∼30 minutes

### A.12 Attention Interpretability

The model provides attention weights for interpretability analysis:

#### Extracting attention weights

The `get_attention_weights()` method returns:

- Protein-to-molecule attention: 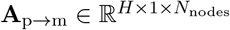
- Molecule-to-protein attention: **A**_m→p_ ∈ ℝ^*H×*1*×*1^ for each of the 2 cross-attention layers.

#### Interpretation

High attention weights indicate which molecular substructures are most relevant for binding to specific protein regions (encoded in the aggregated protein embedding). This can guide fragment-based optimization and SAR analysis.

### A.13 Limitations and Future Directions

#### Current limitations

1. **Training bias**: Model trained on ChEMBL may underperform on extreme chemotypes (PROTACs, macrocycles, peptides)
2. **No 3D information**: Relies on 2D graphs and 1D sequence; misses conformational effects
3. **Uncertainty**: No calibrated confidence intervals on predictions
4. **Single-task**: Does not model off-targets, toxicity, or ADME

#### Planned improvements

1. **3D augmentation**: Incorporate 3D conformer features via distance matrices or geometric GNNs
2. **Uncertainty quantification**: Implement deep ensembles or evidential regression
3. **Multi-task learning**: Joint training on affinity + selectivity + ADME endpoints
4. **Covalent modeling**: Explicit warhead reactivity features and reaction site prediction
5. **Active learning**: Iterative refinement with experimental ABPP data

### A.14 Code and Data Availability

All code, trained models, and processed datasets are available at: https://github.com/ABHCI15/AffinityPred

Includes:

- Complete training and inference scripts
- Model architecture definitions
- Data preprocessing pipelines
- Pretrained model checkpoints
- Screening results for CysDB and MolGlueDB
- Jupyter notebooks for result visualization
- Environment specification files

## References

[1] Jesus P Palafox-Hernandez, Lisa M Boatner, R Gundry, Devin K Schweppe, Keri M Backus, et al. Cysdb: a human cysteine database based on experimental quantitative chemoproteomics. Cell Chemical Biology, 2023. doi: 10.1016/j.chembiol.2023.03.002. URL https://www.cell.com/cell-chemical-biology/fulltext/S2451-9456(23)00109-7.

[2] Xinyu Wang, Ziye Zhuang, Chen Zhang, Xing Fang, Xiaohong Chen, et al. MolGlueDB: an online database of molecular glues. Nucleic Acids Research, 53(D1):D1231–D1238, 2025. doi: 10.1093/nar/gkae882. URL https://academic.oup.com/nar/article/53/D1/D1231/7902517.

[3] Hakime Öztürk, Arzucan Özgür, and Ethem Ozkırımlı. DeepDTA: deep drug–target binding affinity prediction. Bioinformatics, 34(17):i821–i829, 2018. doi: 10.1093/bioinformatics/bty593. URL https://academic.oup.com/bioinformatics/article/34/17/i821/5093245.

[4] Thin Nguyen, Ha Le, Thomas P Quinn, Tri Nguyen, Trung Dung Le, and Svetha Venkatesh. GraphDTA: predicting drug–target binding affinity with graph neural networks. Bioinformatics, 37(8):1140–1147, 2021. doi: 10.1093/bioinformatics/btaa921. URL https://academic.oup.com/bioinformatics/article/37/8/1140/5907191.

[5] Yang Zhang, Caiqi Liu, Mujiexin Liu, Tianyuan Liu, Hao Lin, Cheng-Bing Huang, and Lin Ning. Attention is all you need: utilizing attention in AI-enabled drug discovery. Briefings in Bioinformatics, 25(1):bbad467, November 2023. ISSN 1477-4054. doi: 10.1093/bib/bbad467.

[6] Zeming Lin, Hannes Akin, Roshan Rao, Brian Hie, Siddharth Goyal, and et al. Evolutionaryscale modeling for atomic-level protein structure prediction with a language model. Science, 379(6637):1123–1130, 2023. doi: 10.1126/science.ade2574. URL https://www.science.org/doi/10.1126/science.ade2574.

[7] Saro Passaro, Gabriele Corso, Jeremy Wohlwend, Mateo Reveiz, Stephan Thaler, Vignesh Ram Somnath, Noah Getz, Tally Portnoi, Julien Roy, Hannes Stark, David Kwabi-Addo, Dominique Beaini, Tommi Jaakkola, and Regina Barzilay. Boltz-2: Towards Accurate and Efficient Binding Affinity Prediction, June 2025. URL http://biorxiv.org/lookup/doi/10.1101/2025.06.14.659707.

[8] Saro Passaro, Gabriele Corso, Jeremy Wohlwend, Mateo Reveiz, Stephan Thaler, Vignesh Ram Somnath, Noah Getz, Tally Portnoi, Julien Roy, Hannes Stark, David Kwabi-Addo, Dominique Beaini, Tommi Jaakkola, and Regina Barzilay. Boltz-2: Towards accurate and efficient binding affinity prediction. bioRxiv, 2025. doi: 10.1101/2025.06.14.659707. URL https://www.biorxiv.org/content/10.1101/2025.06.14.659707v1.

[9] Yuyang Wang, Jiarui Lu, Navdeep Jaitly, Joshua M. Susskind, and Miguel Ángel Bautista. Simplefold: Folding proteins is simpler than you think. arXiv preprint arXiv:2509.18480, 2025. URL https://arxiv.org/abs/2509.18480.

[10] Anna Gaulton, Louisa J Bellis, A Patrícia Bento, John Chambers, Mark Davies, Anne Hersey, Yvonne Light, Shaun McGlinchey, David Michalovich, Bissan Al-Lazikani, and John P Overington. Chembl: a large-scale bioactivity database for drug discovery. Nucleic Acids Research, 40(D1):D1100–D1107, 2012. doi: 10.1093/nar/gkr777. URL https://academic.oup.com/nar/article/40/D1/D1100/2903954.

[11] Greg Landrum et al. Rdkit: Open-source cheminformatics software. https://www.rdkit.org, 2006. Accessed 2025-10-01.

[12] Minjie Wang, D. Zheng, Zihao Ye, Quan Gan, Mufei Li, Xiang Song, Jinjing Zhou, Chao Ma, Lingfan Yu, Yu Gai, Tianjun Xiao, Tong He, George Karypis, Jinyang Li, and Zheng Zhang. Deep Graph Library: A Graph-Centric, Highly-Performant Package for Graph Neural Networks, August 2020. URL http://arxiv.org/abs/1909.01315. 1909.01315 [cs].

[13] José Manuel Barraza-Chavez, Rana A. Barghout, Ricardo Almada-Monter, Benjamin Sanchez-Lengeling, Adrian Jinich, and Radhakrishnan Mahadevan. Graph Data Modeling: Molecules, Proteins, & Chemical Processes. September 2025. doi: 10.1021/acsinfocus.7e9017. URL https://pubs.acs.org/doi/book/10.1021/acsinfocus.7e9017.

[14] Keyulu Xu, Weihua Hu, Jure Leskovec, and Stefanie Jegelka. How powerful are graph neural networks? In International Conference on Learning Representations (ICLR), 2019. URL https://openreview.net/forum?id=ryGs6iA5Km.

[15] Zhenxing Wu, Jike Wang, Hongyan Du, Dejun Jiang, Yu Kang, Dan Li, Peichen Pan, Yafeng Deng, Dongsheng Cao, Chang-Yu Hsieh, and Tingjun Hou. Chemistry-intuitive explanation of graph neural networks for molecular property prediction with substructure masking. Nature Communications, 14(1):2585, May 2023. ISSN 2041-1723. doi: 10.1038/s41467-023-38192-3. URL https://www.nature.com/articles/s41467-023-38192-3.

[16] ESM Team. ESM Cambrian: Revealing the mysteries of proteins with unsupervised learning. EvolutionaryScale Blog, 2024. URL https://evolutionaryscale.ai/blog/esm-cambrian. mDecember 4, 2024. GitHub: https://github.com/evolutionaryscale/esm.

[17] Jie Zhou, Ganqu Cui, Shengding Zhang, Cheng Yang, Zhiyuan Liu, Lifeng Wang, Changcheng Li, and Maosong Sun. Graph neural networks: A review of methods and applications. AI Open, 1:57–81, 2020. doi: 10.1016/j.aiopen.2021.01.001.

